# Tissue Engineered Vascularized Patient-Specific Temporomandibular Joint Reconstruction in a Yucatan Pig Model

**DOI:** 10.1101/2020.01.09.900746

**Authors:** Shelly Abramowicz, Sarah Jo Crotts, Scott J. Hollister, Steve Goudy

## Abstract

**Purpose:** Pediatric temporomandibular joint (TMJ) reconstruction occurs as a result of acquired, developmental, and/or congenital conditions. Current pediatric reconstruction options are limited. The aim of this project was to develop a proof of principle porcine model for a load bearing, customized 3-dimensional (3D) printed and BMP2-coated scaffold implanted in a pedicled (temporal) flap as a regenerative approach to pediatric TMJ mandibular condyle reconstruction.

**Materials and Methods:** Scaffolds were custom designed and 3D printed based on porcine computed tomography and absorbed with BMP2. Two operations occured: (1) implantation of scaffold in temporalis muscle to establish vascularity, and six weeks later, (2) unilateral condylectomy and rotation of the vascularized scaffold (with preservation of superficial temporal artery) onto mandibular ramus defect and secured with titanium screws. At 6 months post-implantation, the pigs were sacrified. The experimental side (muscle-scaffold) and the control side (unoperated condyle) were individually harvested at 6 months and evaluated by clinical, mechanical, radiographic, and qualitative/histologic methods.

**Results:** Scaffolds maintained physical properties similar in appearance to unoperated condyles. The vascularized scaffolds had bone formation at edges and adjacent to scaffold-bone interface. New bone was visible in scaffold. Condyle height on the reconstructed side was 68% and 78% of the control side. Reconstructed condyle stiffness was between 20% and 45% of the control side.

**Conclusion:** In our porcine model, customized 3D printed TMJ scaffolds impregnanted with BMP2 and implanted in and pedicled on temporalis muscle has the ability to (1) reconstruct a TMJ defect model, (2) maintain appropriate condylar height and upper airway diameter, and (3) generate new bone, without impacting functional outcomes.

## INTRODUCTION

The pediatric temporomandibular joint (TMJ) may require reconstruction as a result of acquired (eg juvenile idiopathic arthritis, JIA (Abram), trauma), or congenital (hemifacial microsomia, HFM) (Kaban 1988) conditions[1, 2]. The condyle is the primary growth center of the mandible; skeletal growth and development of the condyle are necessary for appropriate function and esthetics. Current options for pediatric TMJ reconstruction are usually only considered in late adolescence and include distraction osteogenesis (DO), autologous reconstruction [i.e. costochondral graft (CCG) or free fibula bone flap (FFF)], and total alloplastic TMJ replacement (TJR). DO requires a long activation period, can fail with premature bone consolidation and may result in poor occlusion and asymmetry[3, 4]. DO may not be applicable in cases where the condyle ramus unit (RCU) is missing (eg HFM). Autologous reconstruction with CCG depends on surrounding recipient tissues for revascularization. Postoperatively, some resorption of CCG will likely occur with potential for complete resorption. FFF usually does not resorb, but the donor defects can be significant. FFF may not routinely be considered in young children due to small caliber of vessels[5, 6] and technical challenges. TJR is not often considered in children because of remaining craniofacial growth potential and likelihood that they would have to be replaced at least once during the child’s lifetime[7, 8]. Due to the current reconstructive limitations, a better reconstructive solution is needed for children who are missing their TMJ.

Developing a regenerative solution that harnesses the body’s ability to form bone and utilizes a local blood supply, without significant donor defect, would be ideal. In early engineering experiments, pre-fabricated TMJ construct implantation with free flaps have been attempted[9], but without reliable vascularization. The ability to rotate a local vascularized flap into a TMJ defect eliminates the complexity of implanting a free flap. We have previously accomplished the design, 3-dimensional (3D) printing, and bone morphogenic protein (BMP)2 delivery necessary to create customized TMJ scaffold/biologic constructs in both large (Yucatan minipig and Yorkshire domestic pig) and small animal models[10-13] Specifically, we: (1) designed and optimized scaffolds to meet patient specific anatomy and functional requirements[10, 13], (2) printed a 3-dimentional customized scaffold designs using polycaprolactone (PCL)[14], (3) delivered BMP2 and BMP2/EPO combinations[11, 12], and (4) demonstrated bone and vascular growth in large and small animal models[10-13]. However, this technology has not been used to reconstruct a pediatric TMJ.

The aim of this project was to develop a large animal model for a load bearing, customized, 3D printed scaffold with BMP-2 and embedded in a vascularized pedicled (temporal) flap as a regenerative approach to pediatric TMJ mandibular condyle reconstruction.

## MATERIALS AND METHODS

### Animal Model

Three male Yucatan minipigs (age 6 months) were used during this experiment. The study was performed in accordance with the regulations and approval of the Institutional Animal Care and Use Committee (internal study code# GT39P) of the T3 Laboratories Inc. in Atlanta, GA and Georgia Institute of Technology. The animals were housed for five days prior to surgery to become acclimated to housing and diet. Throughout the experiment duration, they were monitored daily for general appearance, biologic functions (eg food and water intake and elimination, weight, and activity) and when applicable, signs of pain.

This experiment consisted of two operations which took place under general anesthesia. During the first operation, the scaffold was implanted in the temporalis muscle in order to become vascularized by the temporalis artery. Animals were then allowed to eat, function, and grow normally. Six weeks later, the second operation occurred. Via a hemicoronal incision, the tempoparietal pedicle-scaffold was identified via doppler of temporalis muscle. A unilateral (left side) condylectomy took place and the vascularized scaffold (with preservation of superficial temporal artery) was rotated inferiorly 180 degrees into mandibular ramus and secured with screws (KLS Martin). At end of experiment, animals were sacrificed. The experimental side (scaffold-pedicle) and the control side (unoperated condyle) were individually analyzed (Figure 1).

**Figure 1:**
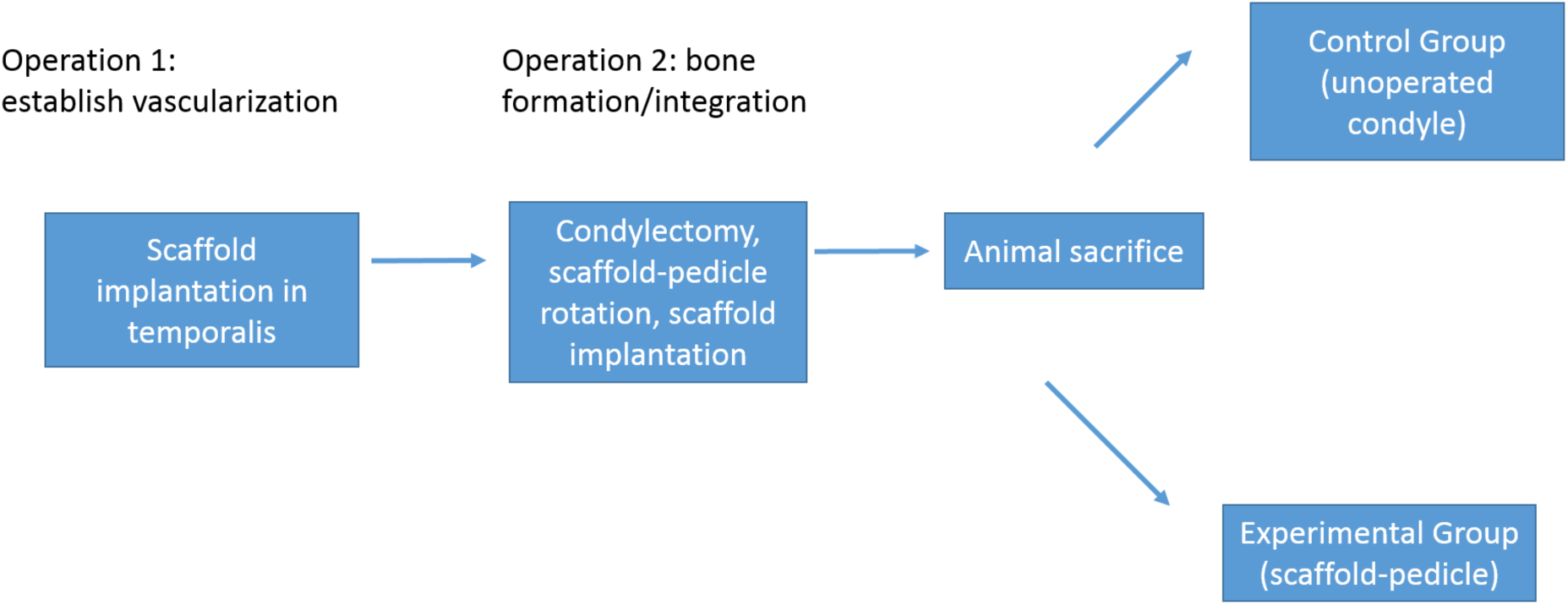
sequence of events.

### Scaffold construction

A customized (pig)-specific porous scaffold of a mandibular condyle was designed directly from a computed tomography (CT) scan of a 6-month old Yucatan minipigs. A Schwarz P triply periodic minimal surface (TPMS) of 40% volume fraction was generated within the mandibular condyle anatomy using image-based design methods[10]. The Schwarz P TPMS microstructure was chosen as it is known to provide an optimal balance between mechanical stiffness and mass transport to support load bearing and tissue growth[15]. An integrated collar with screw holes was created to secure the scaffold to the remaining ramus with titanium screws. The resulting image-based condyle design was converted into STL format using Materialise Mimics^™^ (www.materialise.com). The mandibular condyle scaffold was 3D printed from polycaprolactone (PCL), a resorbable biopolymer, using a laser sintering approach previously described[10, 14, 16, 17]. The resulting scaffold was fit to a 3D Printed model of a Yucatan Minipig mandible (Figure 2).

**Figure 2:**
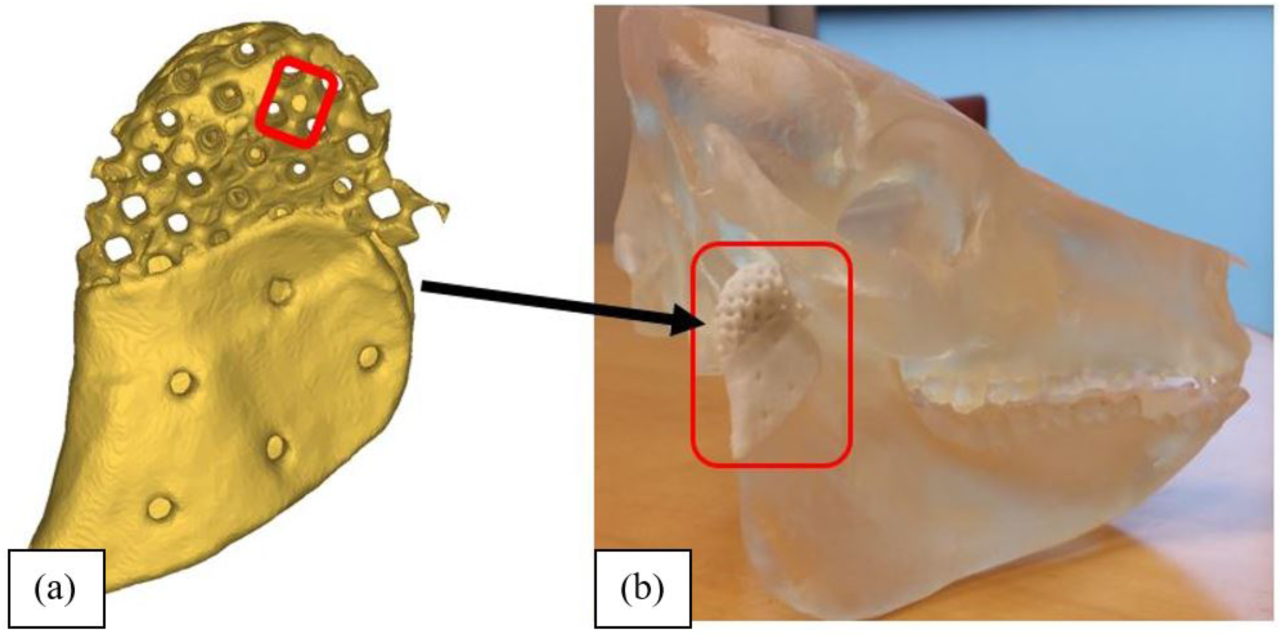
Mandibular condyle scaffold with collar fixation. (a) Design showing repeated Schwartz P microstructure outlined in red (b) 3D printed PCL porous scaffold made of Schwartz P TPMS microstructure with collar fixation fit (outlined in red) on 3D printed Yucatan mandible

### Scaffold insertion

Scaffolds were sterilized in ethylene oxide, then soaked in a solution of 1mg BMP-2 in the operating room. During the first operation, the scaffold was implanted in the temporalis muscle under the superficial temporal artery in order to become vascularized (Figure 3a). After 6 weeks *in situ* (in muscle), the scaffold (now vascularized within the temperoparietal flap) was rotated into the created ramus defect and attached to the ramus with titanium screws (Figure 3b). The untreated condyle served as a control. Animals were then allowed to eat, function, and grow normally for 6 months. Animals were then sacrificed.

**Figure 3:**
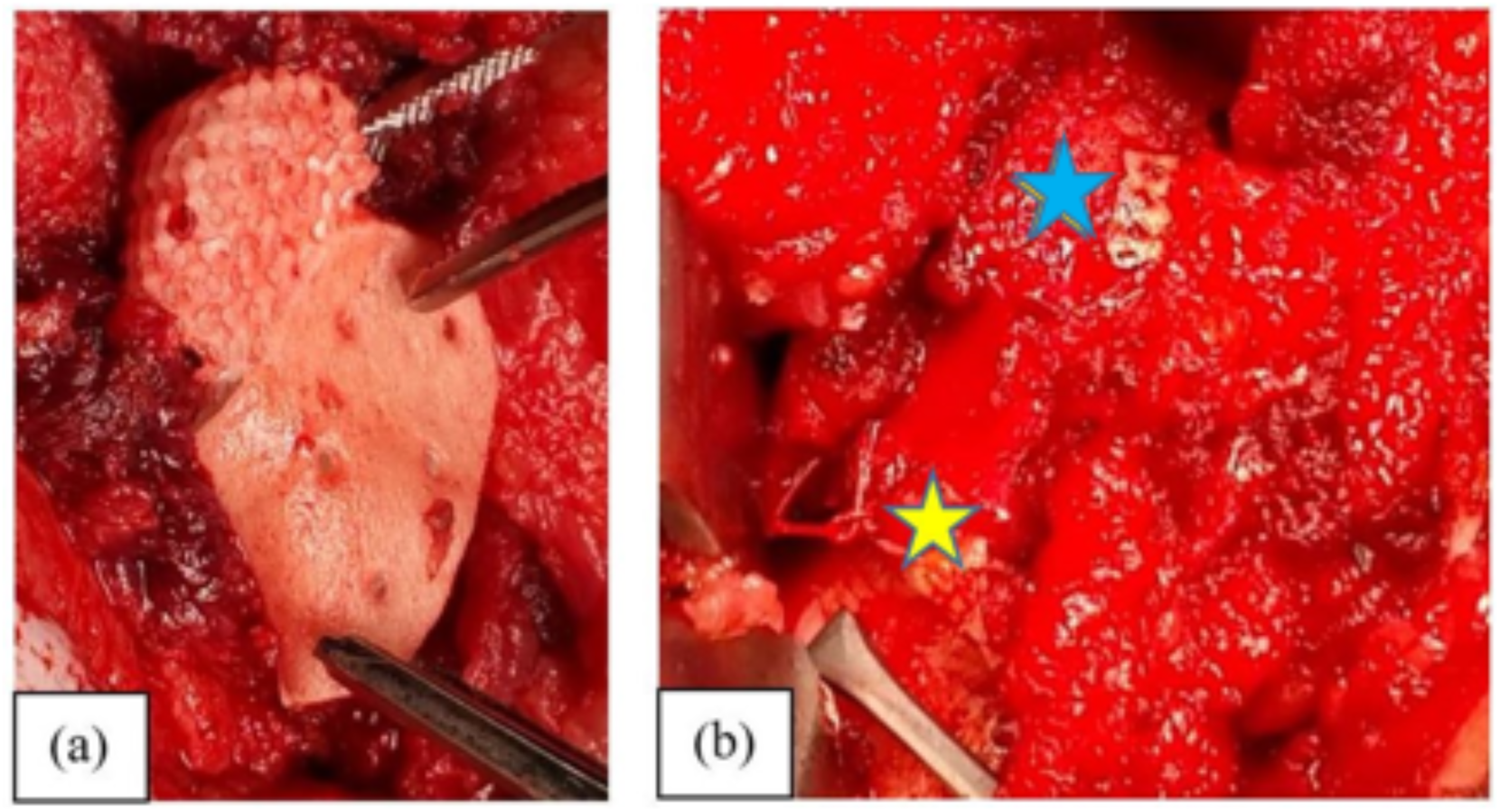
Two stage surgery. Initial scaffold implantation in temporalis muscle (a). Scaffold flap construct (blue star) after rotation to ramus (yellow star) (b).

### Radiologic Analysis

Six months after condyle reconstruction, animals were sacrificed and computer tomography (CT) scans were taken. MicroCT scans and mechanical compression tests were then performed on reconstructed (experimental) condyle and contralateral (non-operated, e.g. control) condyles. CT scans were used to compare new condyle to control condyle by calculating condyle height, condyle volume, and condyle bone volume fraction (total bone volume/condyle volume). Retrieved regenerated scaffolds with new condyle and retrieved contralateral mandibular condyles were tested in mechanical compression along the medial-lateral condyle polls (Figure 4). Resulting load-displacement curves were used to calculate geometric stiffness (load/displacement) from the linear portions of the load-displacement curves. The resected condyle represents condyle stiffness for a Yucatan pig at 7 months of age (6 months old plus 1 month from flap implantation to flap rotation). The contralateral condyle represents condyle stiffess for the Yucatan pig at 13 months of age (6 months old at time of flap implantation, 1 month for flap growth, 6 months until end of experiment).

**Figure 4.**
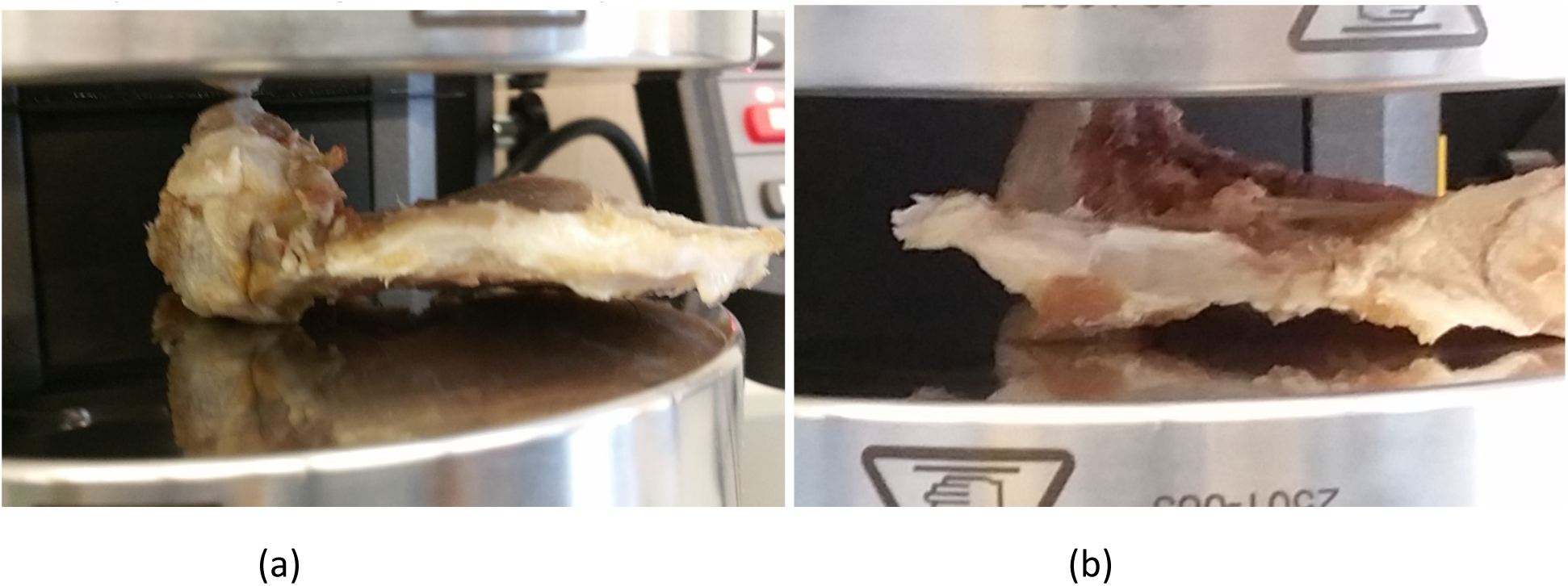
Example of mechanical compression testing (a) Compression along medial-lateral condyle pole of Pig 2. Condyle head, including rotated muscle flap, while remaining mandibular ramus is to the right of the picture. (b) Pig 2 contralateral condyle (non-operated side), tested in compression along medial-lateral pole.

### Histological Analysis

Each specimen was trimmed and embedded in methyl methacrylate. One ground section and three thins sections were taken from the approximate center of the specimen (Figure 5). They were stained (toluidine blue, hematoxylin&eosin, and Safranin O). All sections were evaluated microscopically to determine: (1) bone formation within the construct, (2)regeneration of cartilage at the articulating surface, (3) vascularization, and (4) cellular response to the implants. Photomicrographs were taken as needed to illustrate the findings.

**Figure 5:**
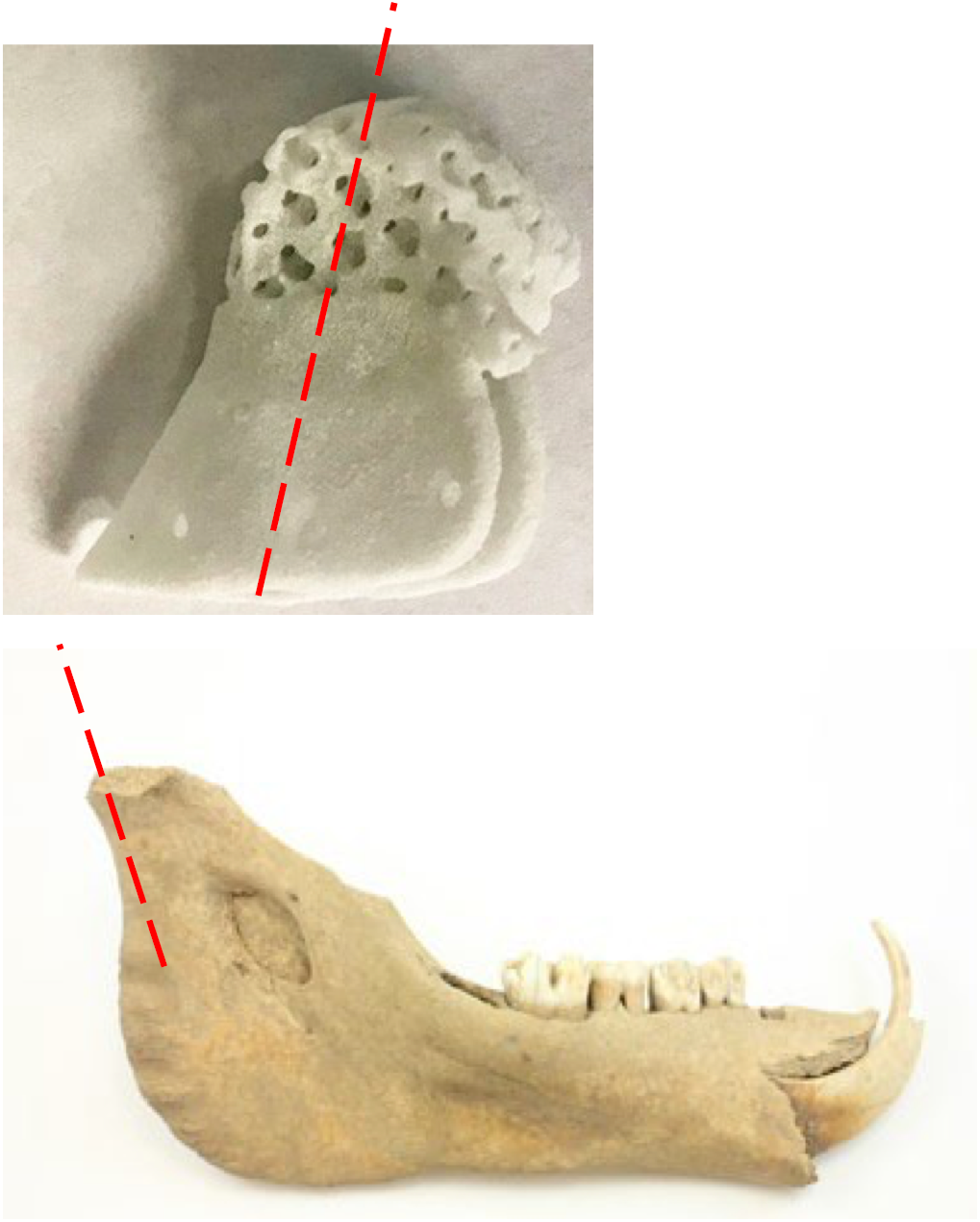
The red dashed lines indicate the plane of section through the implant/condyle.

## RESULTS

### Clinical Evaluation

One animal developed an infection at the implant site due to thin skin adjacent to the surgical site and had to be euthanized prior to conclusion of study. Two animals were allowed to survive for 6 months (weight 32.4kg and 29.6kg). They demonstrated normal bodily functions (mastication, food and water intake and elimination), activities, and weight gain (11.8kg and 13.4kg) during that time.

### Radiologic Evaluation

Models reconstructed from the CT scans demonstrated bony regrowth of the mandibular condyle that had been resected (Fig. 6). Reconstructed condyles had 78% (Figure 6a) and 68% (Figure 6b) of the height of contralateral control condyles. From microCT scans, reconstructed condyle has 67% and 36% of the total condylar volume of the contralateral control condyles. Finally, the bone volume fraction (total bone volume/condyle volume) for the reconstructed condyles were 107% and 105% of the contralateral control condyle bone volume fraction.

**Figure 6:**
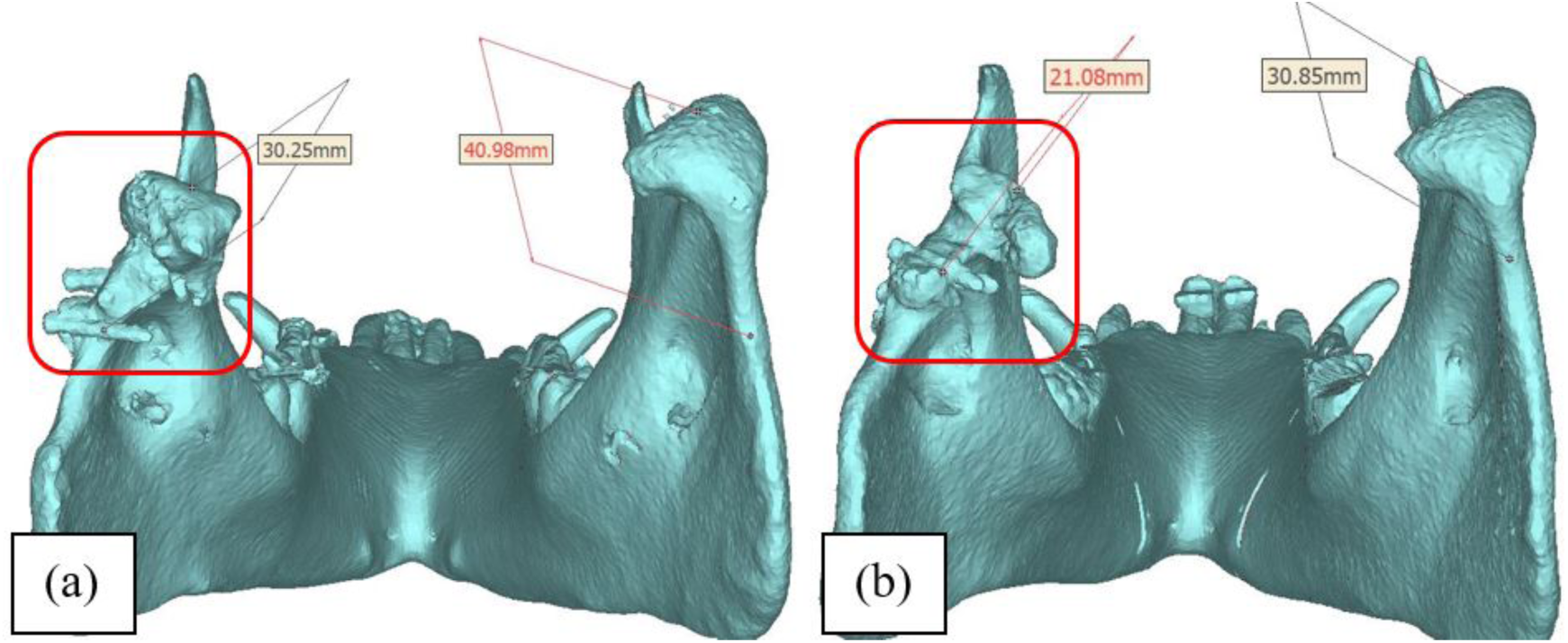
3D rendering of mandible CT scans taken 6 months after flap rotation and condyle reconstruction. Reconstructed side is outlined in red (a) Animal 1 showing reconstructed and contralateral condyle height. (b) Animal 2 showing reconstructed and contralateral condyle height.

### Scaffold Mechanical Evaluation

Load-displacement curves (n=5, 2 regenerated condyles, 2 contralateral/non-operated condyles, and 1 condyle resected at the time of flap rotation) show that the regenerated condyles had lower geometric stiffness than the contralateral or resected condyle (Fig. 7). Regenerated condyles had 36% to 77% of the stiffness of the resected condyles and 20.6% to 46% of the stiffness of the contralateral condyles. The resected condyle stiffness had 57% of the contralateral condyle stiffness. The toe region at the beginning of the load displacement curve results from compression of soft tissue surrounding the bony condyle.

**Figure 7:**
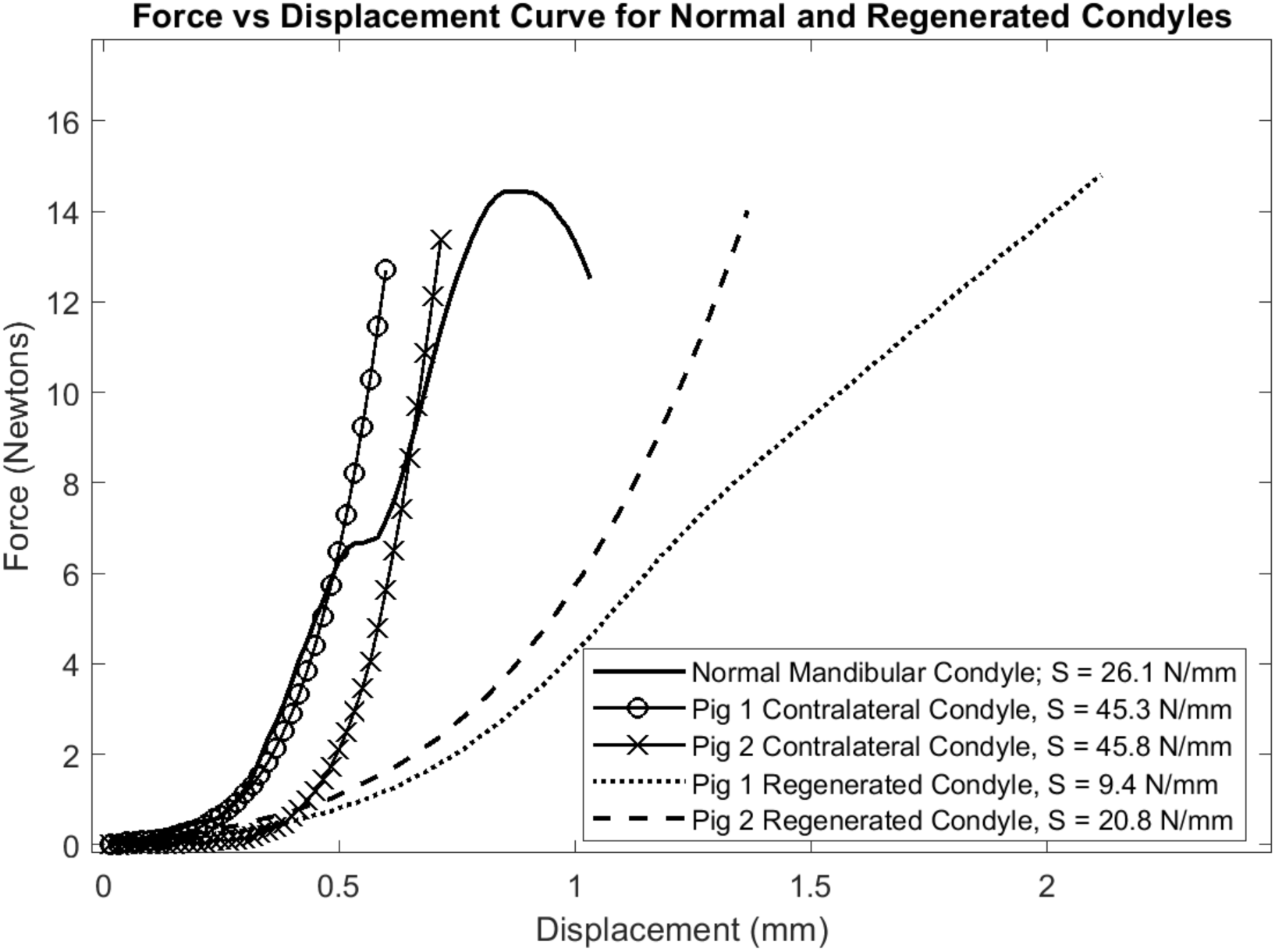
Load-displacement curves for compression testing of regenerated and resected (control) condyles. Linear portion of load-displacement curve represents stiffness of bony region. Regenerated condyles had 36% to 77% of the stiffness (S) of the resected condyles and 20.6% to 46% of the stiffness of the contralateral condyles. The resected condyle stiffness had 57% of the contralateral condyle stiffness.

### Qualitative Analysis/Histological Evaluation

Thin sections from the approximate center of the neo-condyles, contralateral controls, and mandibular condyle scaffolds were evaluated microscopically. The control and experimental condyles were compared. The results demonstrated that there was new bone in a few focal within the implant (figure 8) and that the implant was partially incorporated in newly formed bone (figure 9). There was no cartilage regeneration. Blood vessels were visualized at surface of implant material (fig 10). Vascularized fibrous tissue (fig 11) with macrophage and giant cell infiltrates were found to be associated with the implant material.

**Figure 8:**
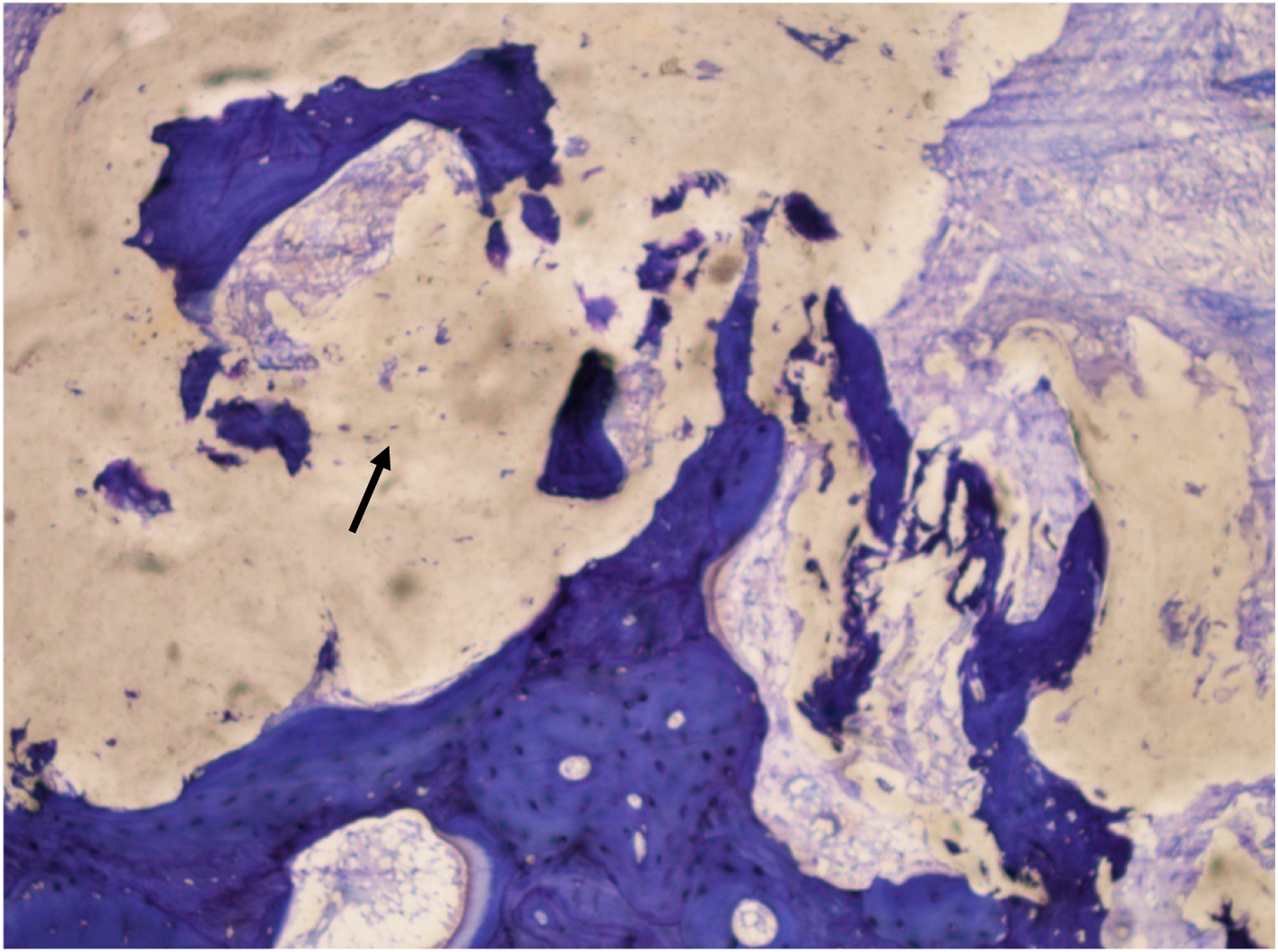
Bone infiltration into the implant (arrow). Toluidine Blue; original magnification, 100X.

**Figure 9:**
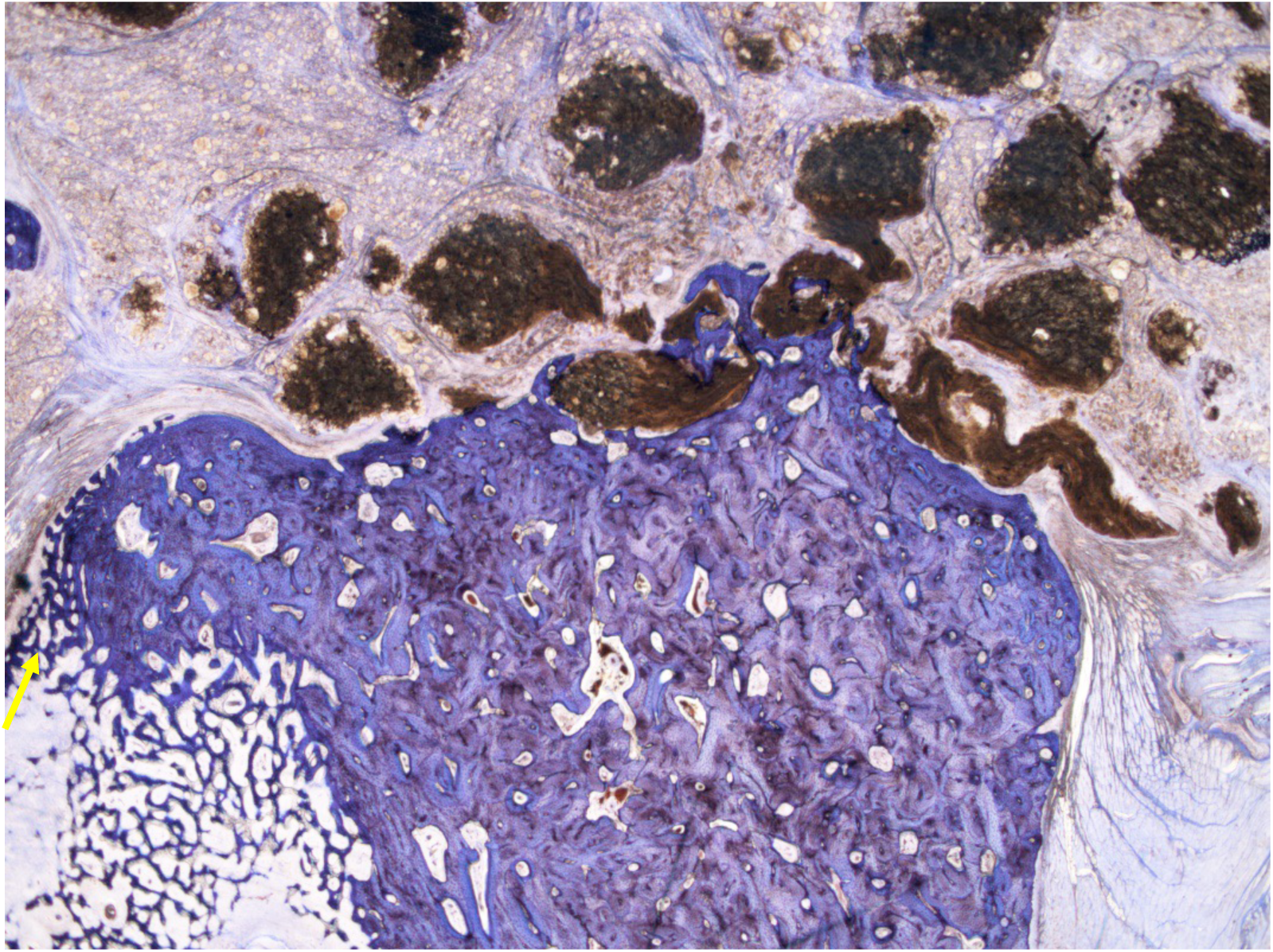
The implant is partially incorporated in newly-formed bone (woven bone, arrow). Toluidine Blue; original magnification, 10X.

**Figure 10:**
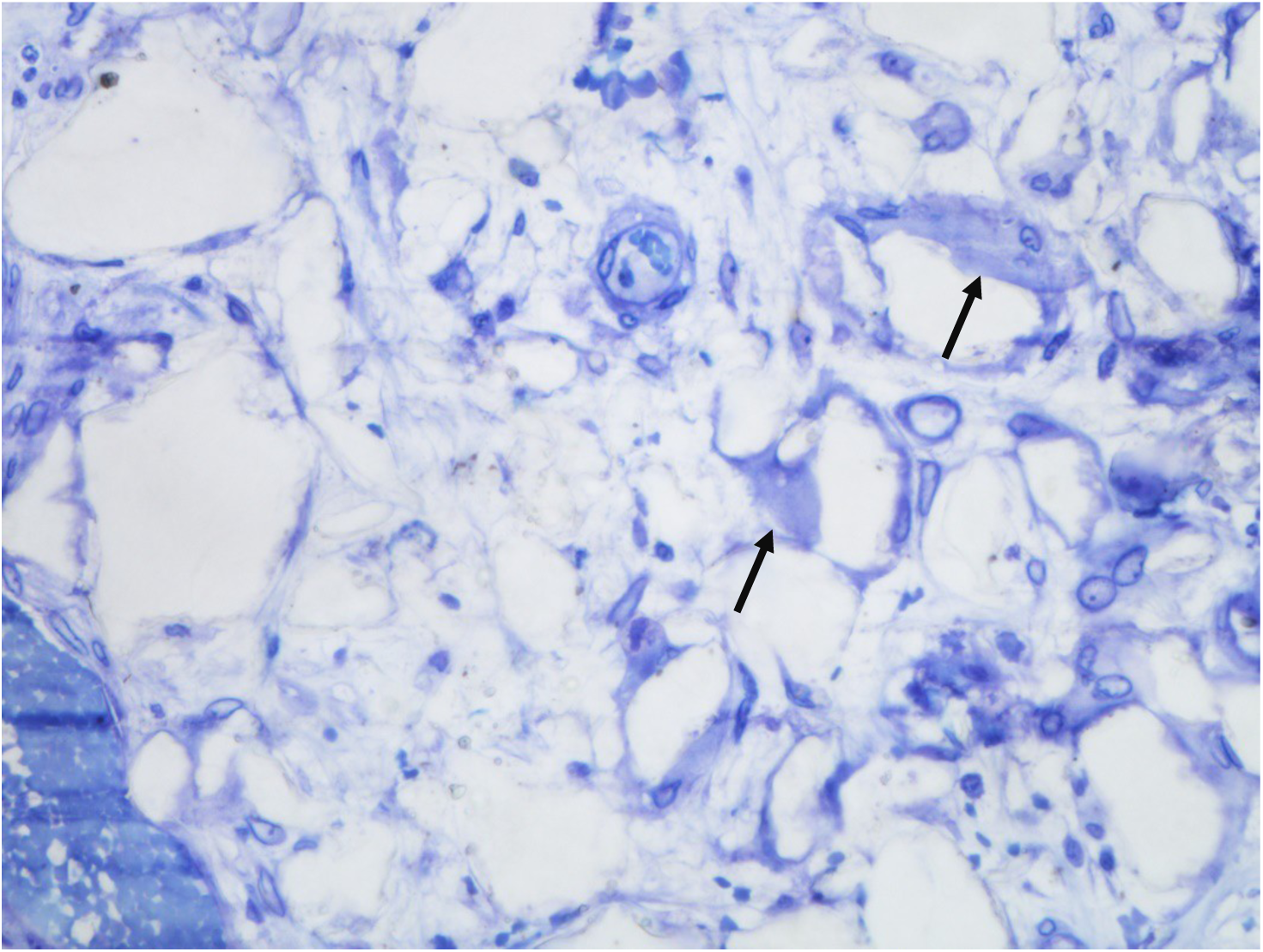
Multiple blood vessels with giant cells (arrows) at surface of implant material (seen as voids). Toluidine Blue; original magnification, 200X

**Figure 11:**
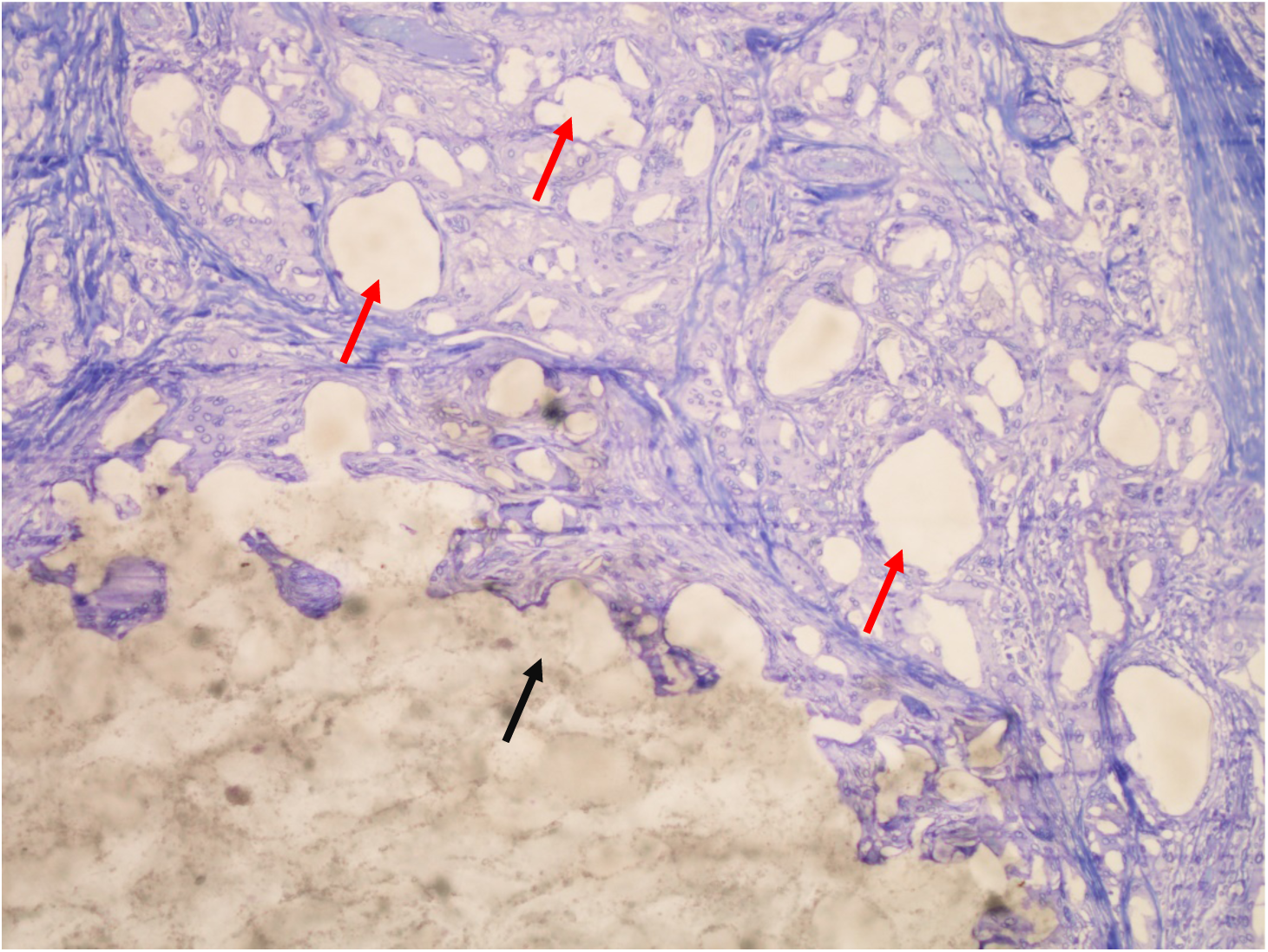
Vascularized fibrous tissue can be seen between the bulk material (black arrow) and smaller pieces of implant material (red arrows) seen as voids (section 18-033-6A; animal 59-146L). Toluidine Blue; original magnification, 100X.

## DISCUSSION

The aim of this project was to develop a model for a load bearing, customized (porcine), 3D printed and expandable scaffold, absorbed with BMP-2 based on a pre-vascularized pedicled (temporal) flap as a regenerative approach to pediatric TMJ mandibular condyle reconstruction.

In our project, we compared CT scan of the reconstructed condyles to the un-operated (control) TMJ. This revealed that the reconstructed 3D printed PCL TMJ BMP2 maintained mandibular height, condylar volume, and bone volume fraction. These results are similar to outcomes reported by Bhumiratana et al.[18] who reported pre-seeding decellurarized bovine bone condyle scaffolds with 100 to 150 million adipose derived stem cells for 3 weeks prior to implantation. This technology, consisting of cell therapy outside of cells derived from the patient in the operating room, faces a significant regulatory hurdle that would make it difficult to translate such an approach for pediatric mandibular condyle reconstruction. However, the ability to design and fabricate a custom patient-specific scaffold via a custom device exemption or expanded access may enable this approach to be clinically translated for a pediatric patient population.

In our study, we used 3D printing technology to form the scaffold of condyles. This technology is being used in many facets of pediatric and adult regenerative medical therapies (i.e. ear and airway reconstruction). Previous studies on 3D tissue regeneration are performed on small animal models, but few use large animal models that mimic human tissue regeneration. Limitations in 3D printed constructs exist with scaffold vascularization[19] and integration by the patient. By combining 3D printing technology with BMP2 bone regeneration technology, we showed that BMP2 impregnated TMJ 3D printed scaffolds are able to regenerate a functional TMJ with quantifiable bone measurements using a vascularized 2-step temporalis flap approach.

Regenerative solutions to bone loss in adults have already been accomplished in spine and maxillary surgery using BMP2[20, 21]. However, a regenerative solution in children is needed. An ideal TMJ reconstruction would allow for stable biological properties necessary for daily function while bone regeneation is occuring. Using our protocol, the functional outcomes of eating and weight gain were maintained by the pigs. We demonstrated that the reconstructed TMJ was able to preserve height and volume necessary for daily activities. Functionally, weight or diet did not change following surgery, showing that the surgical intervention did not interfere with regular activites of daily living.

The integration of the temporalis muscular flap to vascularize the regenerative scaffold is based on the the superficial temporal artery blood supply. This method is currently routinely used for head and neck reconstruction [22](Dallan). Most commonly, this flap is used to cover auricular prostheses that are implanted to address microtia[23] (yang 2009). Previous models stated that development of vascularization has been challenging[19, 24] (Abukawa 2003, 2004). Using our juvenile mini-pig TMJ condylectomy model, 3D printed osteoinductive PCL scaffold impregnated with BMP2 was used in a two step manner. Our study demonstrated partial cell and fibrous tissue infiltration of the constructs implanted in the temporalis muscle. The tissue present was predominantly vascularized fibrous tissue, with macrophage and giant cell infiltrates associated with the implant material. Blood vessels were visualized at surface of implant material demonstrating angiogenesis and recruitment of blood vessels necessary for bone formation. These results suggest that the implants were likely vascularized. Thus, our protocol of using a pedicled blood supply, proved to be efficacious. Lastly, our protocol demonstrated new bone formation in the neo-condyles. This bone served as initial matrix for additional bone formation and allowed normal mandibular function. Additional large animal studies will further delineate the durability and long-term (>1 year) bone regeneration and functional outcomes. Current limitations in this pilot study is the loss of one animal and the 6 month time point, whereas additional bone ingrowth is expected to occur as the PCL scaffold slowly resorbs over a several year time frame. A challenge for pre-fabricated flap approaches going forward will be delivering appropriate bone and vascular biologic signals to develop bone and vascular tissue in the flap prior to rotation. Addressing this challenge will require experimentation using a variety of biologics and delivery vehicles within the load bearing scaffolds.

In summary, we demonstrated that in our porcine model, temporalis flap-based 3D printed PCL TMJ with BMP2 has the ability to (1) reconstruct a TMJ defect model, (2) maintain appropriate condylar height and (3) generate new bone without impacting functional outcomes. Bone formation in the flap prior to rotation was minimal. We are exploring other BMP2 delivery methods from the 3D printed scaffold, including a PolyEthyleneGlycol (PEG) based hydrogel system to allow controlled BMP2 delivery. Futhermore, the scaffold structure could be optimized to enhance its load bearing capability.

## ACKNOWLEDGEMENTS

This study was supported by a pilot grant from the Regenerative Engineering and Medicine center, a joint collaboration between Emory University and Georgia Institute of Technology. We would like to thank Mr. Ryan Akman for assistance with mechanical testing and microCT scanning as well as Dr. Harsha Ramaraju for assistance with the BMP2 adsorption on the scaffold. T3 Labs (gcmiatl.com/preclinical-testing-and-training) for animal surgical support and animal CT scanning.

